# Historical causes for the greater proportion of polyploid plants in higher latitudes

**DOI:** 10.1101/2023.09.01.555981

**Authors:** Eric R. Hagen, Thais Vasconcelos, James D. Boyko, Jeremy M. Beaulieu

## Abstract

**Premise of the Study:** The proportion of polyploid plants in a community increases with latitude, and different hypotheses have been proposed about which factors drive this pattern. Here, we aim to understand the historical causes of the latitudinal polyploidy gradient using a combination of ancestral state reconstruction methods. Specifically, we assess whether (1) polyploidization enables movement to higher latitudes (i.e., polyploidization precedes occurrences in higher latitudes) or (2) higher latitudes facilitate polyploidization (i.e., occurrence in higher latitudes precedes polyploidization).

**Methods:** We reconstruct the ploidy states and ancestral niches of 1,032 angiosperm species at four paleoclimatic time slices ranging from 3.3 million years ago to the present, comprising taxa from four well-represented clades: Onagraceae, Primulaceae, *Solanum* (Solanaceae), and Pooideae (Poaceae). We use ancestral niche reconstruction models alongside a customized discrete character evolution model to allow reconstruction of states at specific time slices. Patterns of latitudinal movement are reconstructed and compared in relation to inferred ploidy shifts.

**Key Results:** We find that no single hypothesis applies equally well across all analyzed clades. While significant differences in median latitudinal occurrence were detected in the largest clade, Pooideae, no significant differences were detected in latitudinal movement in any clade.

**Conclusions:** Our preliminary study is the first to attempt to connect ploidy changes to continuous latitudinal movement, but we cannot favor one hypothesis over another. Given that patterns seem to be clade-specific, a larger number of clades must be analyzed in future studies for generalities to be drawn.

## INTRODUCTION

Polyploidy – i.e., the state of having more than two complete sets of chromosomes – has continually shaped the evolutionary history of flowering plants. Indeed, whole genome duplications are identified along the stem leading to all flowering plants as well as many events occurring all throughout some of the most diverse, as well as some of the most depauperate, clades nested within (Jiao et al., 2011). Through comparisons of diploid and polyploid plants, polyploidy appears linked to a variety of evolutionary changes, including novel phenotypic traits (Levin, 1983), ecological relationships (Segraves, 2017), and macroevolutionary patterns (e.g., Mayrose et al., 2011; Soltis et al., 2014). In biogeography, polyploidy is largely studied in the context of latitudinal and elevational gradients, in which polyploids tend to compose larger proportions of the flora at higher latitudes and elevations than at lower ones (Stebbins, 1950; Brochmann et al., 2004; Rice et al., 2019). The so-called “latitudinal polyploidy gradient” (LPG) has long been observed in individual clades (e.g., Löve and Löve, 1943, 1949), and recent studies incorporating large amounts of distribution data across clades have largely confirmed the generality of this pattern (Rice et al., 2019).

Proposed mechanisms responsible for the creation of the LPG can be divided into two categories. First, conditions of poleward environments lead to higher rates of polyploid formation at higher latitudes. Harsh environmental conditions like cold stress are known to induce polyploidy (De Storme and Geelen, 2014; Lohaus and Van de Peer, 2016), and the fragmented ranges of poleward environments could lead to allopolyploid formation via repeated contacts after range expansion (Stebbins, 1985). Second, various adaptations of polyploids lead them to preferentially move into poleward environments at rates higher than those of diploids. Polyploids are believed to have generally greater colonizing ability than diploids due to higher rates of self-compatibility (Bierzychudek, 1985; Barringer, 2007) and phenotypic plasticity (Price et al., 2003; Leitch and Leitch, 2008). Thus, in the time since freezing conditions began to appear at northern latitudes during the Pliocene (Mudelsee and Raymo, 2005), the LPG could have been generating by plant lineages generally moving to higher latitudes after polyploidization events.

These two scenarios, which we call the “centers of origin” and “centers of arrival” hypotheses, respectively, are not mutually exclusive. If polyploid plants are, in fact, generally better adapted to harsh environments, one may expect to see both greater rates of polyploidization near the poles and greater movement into these environments of polyploids that originate elsewhere. It should also be noted that there remains the possibility that the LPG emerges passively. For example, Rice et al. (2019) found that global polyploid distribution is strongly correlated with climate, though they suggest that this is mainly indirect, as polyploids tend to be perennial (Van Drunen and Husband, 2019), herbaceous plants (i.e., chamaephytes) that are low to the ground and able to survive the harsh conditions of poleward environments (Raunkiaer, 1934). However, while present-day climatic variables do correlate with biogeographic patterns, the modern distributions of plants largely result from past climate changes (Normand et al., 2011). Additionally, correlations between specific traits and environmental variables may be shaped more by shared evolutionary history among species sharing those traits rather than functional relationships (Svenning and Skov, 2007; Ma et al., 2016; Sundaram and Leslie, 2021), so phylogenetic information must be considered as well. In any event, teasing apart the evidence for each scenario across flowering plants would provide invaluable clues about the historical causes for the LPG.

Such an investigation comes with particular challenges. The first is the need to incorporate information about historical plant distributions, which is particularly difficult due to the large number of biotic and abiotic factors that can potentially influence a species’ geographic range. For instance, Rice et al. (2019) included paleoclimatic data from the Last Glacial Maximum (LGM; 21 kya) in their analysis, but this was only used in the context of correlating deglaciation extent with ploidy distributions, and they implicitly assumed that ranges remained unchanged to the present. While discrete-area methods of inferring ancestral ranges (e.g., Ree and Smith, 2008; Matzke, 2014) are in wide use, the reconstructions they create are usually coarse and contain few areas. Continuous-area methods requiring paleoclimatic data and shallower timescales are less common (Guillory and Brown, 2021), but these methods are more powerful for inferring latitudinal movement than tracking movement between arbitrarily designated geographic areas. The second difficulty is reconstructing ploidy changes through time. While ploidy can be reconstructed from fossils with preserved cuticle (McElwain and Steinthorsdottir, 2017), fossil data is too sparse for a large-scale study. Instead, one would need to rely on reconstructions using a model of ploidy state transitions over time. One advantage of using such models is the ability to reconstruct not only polyploidization but also diploidization, which is the reorganization of the genome that returns a plant to a diploid (or “diploid-like”) state after whole genome multiplication. Although we have no expectation of how species will move latitudinally following diploidization, it may be illuminating to compare movement between species that polyploidize as opposed to diploidize, as well as stay polyploid or diploid, as a “control” group.

Here, in what we believe is the first attempt to discern the historic causes of the LPG, we analyze the distributions of plants in historical and phylogenetic context to determine how plants in specific clades move across latitudes after ploidy transitions. Specifically, by analyzing the timing of reconstructed ploidy changes and biogeographic movements, we test the generality of the “centers of arrival” hypothesis, in which range movement towards higher latitudes happens most often after polyploidization events (Fig. 1a), and the “centers of origin” hypothesis, in which polyploids form mostly at poleward environments and subsequently stay or move towards the equator (Fig. 1b).

**Figure 1.**
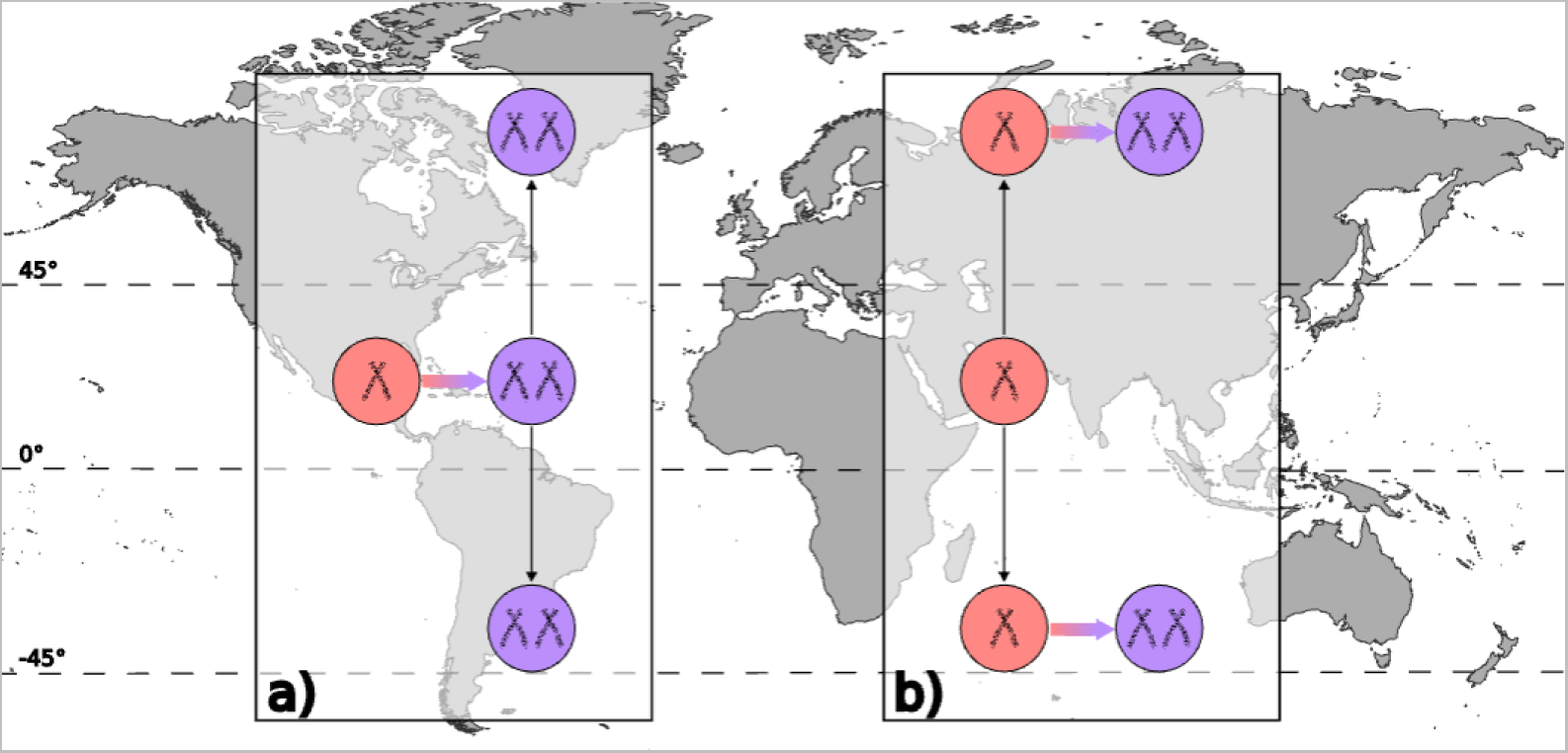
Conceptual diagram of historical biogeographic patterns expected to be observed under the “centers of arrival” hypothesis (a) and the “centers of origin” hypothesis (b). Under the “centers of arrival” scenario, polyploidization occurs across the globe but is followed by higher rates of antiequatorial movement relative to diploids, thus creating the LPG. Under the “centers of origin” scenario, the LPG is created by higher rates of polyploidization in poleward environments.

## MATERIALS AND METHODS

### Phylogenetic and ploidy datasets

We opted to use a multi-clade approach for this work, which aims to discern both biological generalities as well as clade-specific patterns (e.g., Boyko et al., 2023; Vasconcelos, 2023). The main reason for choosing this approach is to reduce the impact of sampling bias in subsequent analyses of ancestral state and ancestral range reconstructions by focusing on clades that are particularly well sampled, as opposed to using super matrix trees (e.g., Smith and Brown, 2018) that are unevenly sampled. These biases are also caused by available ploidy data being skewed toward certain taxonomic groups, particularly those studied in the Global North (Marks et al., 2021), and the fact that available GBIF data are incomplete as well as spatially clustered (Beck et al., 2014).

Our work makes use of four well-represented clades with relatively high availabilities of ploidy data and sampling at the species level: Onagraceae (Freyman and Höhna, 2019), Primulaceae (De Vos et al. 2014), Pooideae (Poaceae; Spriggs et al., 2014), and *Solanum* (Solanaceae; Särkinen et al., 2013). The Onagraceae tree contains 292 species (c. 45% sampling; 186 with ploidy data), Primulaceae contains 263 species (c. 9.4% sampling [Xu and Chang, 2017]; 141 with ploidy data), Pooidae contains 1,312 species (c. 40.6% sampling [Soreng et al., 2017]; 748 with ploidy data), and *Solanum* contains 441 species (c. 33.3% sampling; 256 with ploidy data). The Pooidae and *Solanum* trees were pruned from larger phylogenies of Poaceae and Solanaceae, respectively, because the larger Poaceae and Solanaceae trees had data coverage of less than 50% for ploidy data, and pruning to include only these lower taxonomic rankings allowed us to focus on clades that are particularly data rich. These four clades were selected for this study because they are comparatively large, are well-represented in our ploidy dataset, are geographically widespread, come from different parts of the phylogeny of angiosperms (different major clades – rosids, asterids, and monocots), and because preliminary analyses recovered a relatively large number of recent polyploidization and diploidization events in the Quaternary, our focal time slice.

Ploidy data was extracted from the supplementary data of Rice et al. (2019), which is contained in individual ChromEvol output files separated by genus. We combined these individual files into a master table and filtered it for species represented in our four phylogenies. In our analysis, we define a “polyploid” narrowly to specifically refer to a neopolyploid (i.e., newly formed polyploids; Ramsey and Schemske, 2002), following the methodology of Rice et al. (2019). Neopolyploids are cytologically distinct from their diploid progenitors, and they have undergone whole genome multiplication sufficiently recently that they retain additive genome sizes of their parents as well as distinguishable subgenomes (Mandáková and Lysak, 2018). In contrast, mesopolyploids and paleopolyploids are species that underwent polyploidization further in the past and have undergone diploidization to decrease their genome size as well as genome restructuring. We use this definition for two reasons: (1) the LPG is a gradient of plants that *are* polyploid (i.e., neopolyploids) rather than of plants that *behave* like polyploids (in the sense of gaining advantageous traits rather than chromosomal behavior), and (2) because we examined latitudinal changes after inferred events of both polyploidization and diploidization, so it did not make sense to consider re-diploidized plants in our analysis as polyploids, that is, paleopolyploids (see “Integrating trait evolution models with reconstructions of past climatic niches” below).

### Distribution data

We downloaded all occurrence points available on GBIF that were based on preserved specimens (i.e., excluding human observations) for the four focal clades in our study (GBIF 2022). We then removed inaccuracies following protocols similar to those of Boyko et al. (2023). Our final occurrence point database accounts for 331,434 points in total, including 43,408 for *Solanum*, 11,200 for Primulaceae, 210,461 for Pooideae, and 66,365 for Onagraceae. After filtering for only those phylogenetically represented species with ploidy data and *sufficient* occurrence points (3 or more), the following number of species was analyzed for each clade: 543 in Pooideae, 218 in Solanum, 164 in Onagraceae, and 107 in Primulaceae, for a grand total of 1,032 species.

### Integrating trait evolution models with reconstructions of past climatic niches

Most models that connect biogeographic shifts with discrete trait evolution require modeling areas discretely rather than continuously (e.g., Ree and Smith, 2008; Goldberg et al., 2011; Caetano et al., 2018), and while methods have recently been developed to connect the evolution of discrete traits with continuous ones (e.g., Boyko et al., 2023), modeling range evolution continuously usually involves climatic parameters like temperature and precipitation. Because it is unclear whether the LPG may be caused by climatic factors or other biogeographic causes (e.g., Stebbins, 1985), we opted to instead model range evolution and ploidy evolution separately and test for connections between the two post-hoc.

We began by modeling ploidy shifts along the phylogeny of each clade during the past c. 3.3 million years. To that end, we used corHMM (Beaulieu et al., 2013; Boyko and Beaulieu, 2021) with modified functions that allow for ancestral state reconstruction at specific time slices rather than at nodes. We designed this because we were interested in inferring ploidy states at shared time slices, specifically those with climatic data available from the PaleoClim database (Brown et al., 2018), rather than at asynchronous branching points (i.e., the nodes of a phylogeny) as is the default of the software (see Fig. 2). For each phylogeny, we tested three different model structures, none of which utilized hidden states: ER (equal transition rates between diploid state and polyploid state), ARD (transition rates between diploidy and polyploidy are allowed to vary), and a custom-made unidirectional structure where reversal to diploidy was disallowed after polyploidization. Some models of ploidy evolution (e.g., Robertson et al., 2011) disallow reversals to diploidy based on arguments that ploidy evolution is significantly asymmetrical (e.g., Stebbins, 1971; Meyers and Levin, 2006). However, much research suggests that reversals to diploidy are prevalent in flowering plants (Mandáková and Lysak, 2018), and other models of ploidy evolution reflect this (Zenil-Ferguson et al., 2019). Given our interest in both polyploidization and diploidization, we opted for the latter strategy. We evaluated support for each model using AIC (Akaike, 1974) and a significance criterion of a difference of at least 2 AIC units (Burnham and Anderson, 2002).

**Figure 2.**
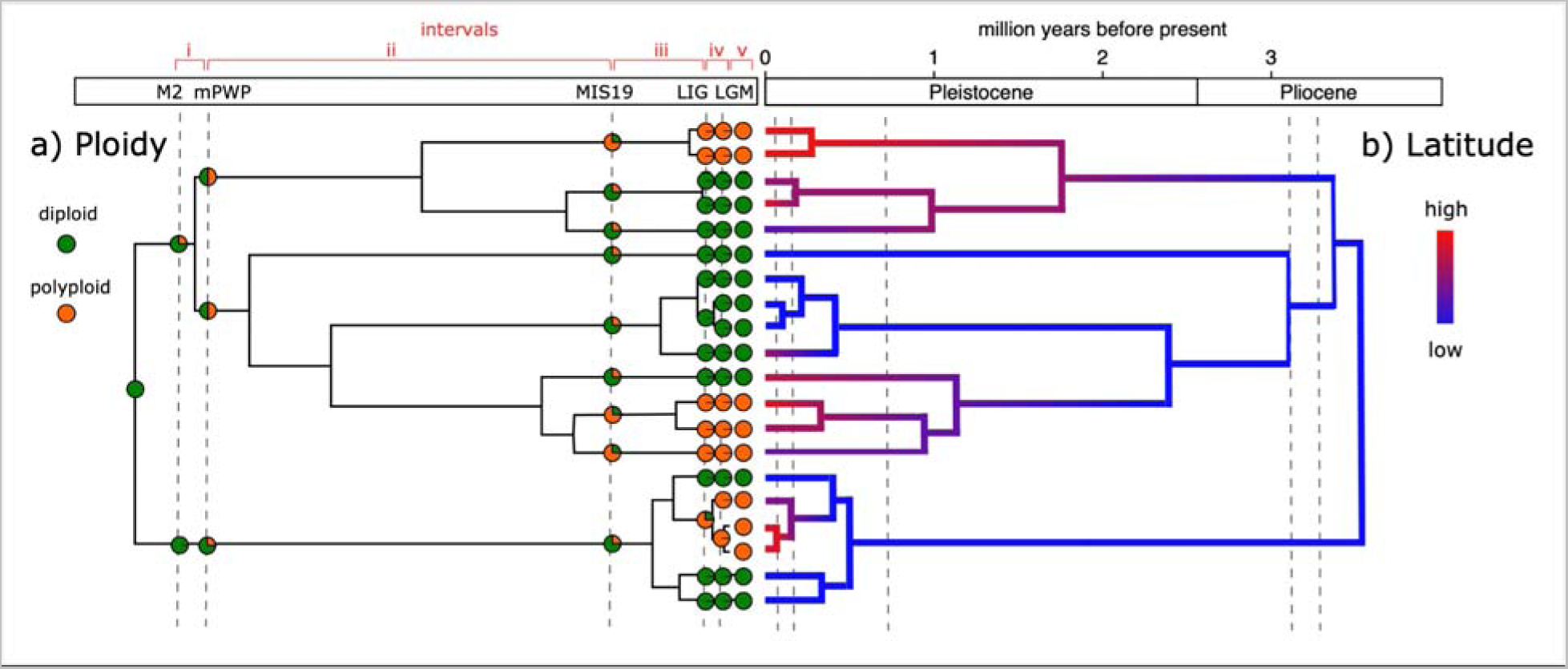
Conceptual figure showing our method of correlating inferred ploidy shifts at paleoclimatic time slices with estimated latitudinal changes, allowing for the connection of ploidy shifts to biogeographic movement. This scenario depicts the expectation under the “centers of arrival” hypothesis, in which shifts in ploidy (a) are followed by antiequatorial latitudinal movement (b).

Once corHMM models were run, we used a novel marginal reconstruction function to calculate the marginal probabilities of anagenetic taxa occurring at the time slices for which we had paleoclimatic data. Once ploidy shifts had been modeled, we reconstructed the range evolution of lineages in each tree using the R package machuruku (Guillory and Brown, 2021), a tool for phylogenetic niche modeling that allows for continuous reconstruction of ranges at time slices with paleoclimatic data as well as visualization of inferred spatial distributions. We reconstructed ranges at 4 time slices based on data from PaleoClim (Brown et al., 2018): the Last Interglacial (LIG, c. 130 ka), Marine Isotope Stage 19 (MIS19, c. 787 ka), the mid-Pliocene Warm Period (mPWP, c. 3.205 Ma), and Marine Isotope Stage M2 (M2, c. 3.3 Ma), all using the spatial resolution of 10 arc-minutes (∼20 km). For each time slice, we first estimated tip response curves to each climatic variable using the function “machu.1.tip.resp,” then estimated the ancestral niches of each taxon extant at each time slice with “machu.2.ace,” and finally projected the ancestral climatic niche models for each slice onto maps containing paleoclimatic variables with “machu.3.anc.niche.” We ran the “machu.3.anc.niche” function with the “clip.Q” option set to False, which produces models including less suitable areas but which prevented the function from returning NA results for some lineages.

### Biogeographic analyses

To examine biogeographic movements through time, we first examined the extent to which the LPG is present among the four separate clades we examined using both ANOVAs and phylogenetic ANOVAs (Revell, 2012). We then parsed latitudinal changes between time slices concurrent with different ploidy transitions, characterizing species ranges by their median latitudes. We divided possible ploidy transitions into four possible ploidy status categories depending on whether ploidy changes happened or not within a given time slice: (1) staying diploid, (2) staying polyploid, (3) diploidization, and (4) polyploidization. Lineages that did not change ploidy (i.e. "staying diploid" and "staying polyploid") are used as null hypotheses against which we can compare species that changed ploidy. The “centers of arrival” hypothesis would be supported when movement towards higher latitudes occurs more frequently after polyploidization than any other category of ploidy change. On the other hand, the “centers of origin” hypothesis will be supported where starting latitudes at the time slice when polyploidization occurs is significantly higher than for the other ploidy change categories. For each category, we tested for significant trends in movement (absolute latitudinal change) using a simple sign test (Conover, 1971), employing the “binom.test” function in R (R Core Team, 2022) to compare median latitudes at the beginning and end of each time slice. To account for the magnitude of change in addition to whether movement was generally equatorial or antiequatorial, we also conducted Wilcoxon signed-rank tests (Wilcoxon, 1945) on the same data, both with and without phylogenetic weights incorporated.

To quantitatively compare whether latitudinal movements across all time slices significantly differed between species that polyploidized and those that diploidized, we conducted two-sided Kolmogorov-Smirnov tests (Smirnov, 1939. Finally, to determine whether species that polyploidize possess ranges at significantly different latitudes relative to species in the other three ploidy status categories, we used phylogenetic ANOVA (Revell, 2012) to compare reconstructed median starting latitudes and latitudinal change across species.

Comparisons were restricted to be conducted within clades and within the same time slices.

## RESULTS

### Model selection and transition rates

Goodness of fit of corHMM models with different assumptions of transitions rates (i.e., equal rates, ER; all rates different, ARD; and unidirectional, uni) between ploidy states within the four clades were mixed (Onagraceae, AIC_ARD_=82.58, AIC_ER_=80.63, AIC_uni_= 105.48; Primulaceae, AIC_ER_ = 87.89, AIC_ARD_=82.96, AIC_uni_=92.74; Solanum, AIC_ER_ = 103.31, AIC_ARD_=103.31, AIC_uni_=103.0; Pooideae, AIC_ER_ = 503.09, AIC_ARD_=498.46, AIC_uni_=586.63); however, the model with best fit in all groups always allowed some transitions between the two states. In other words, the unidirectional model was never favored. Ploidy transitions were reconstructed using ARD in Primulaceae and Pooideae, as it was favored by >2 AIC units, and with ER in Onagraceae and *Solanum*, because we defaulted to the model with fewest parameters since no model was favored by AIC comparison. Inferred rates of polyploidization were 0.019 transitions Myr^-1^ in Onagraceae, 0.09 transitions Myr^-1^ in Primulaceae, 0.02 transitions Myr^-1^ in *Solanum*, and 0.21 transitions Myr^-1^ in Pooideae. Rates of diploidization were 0.019 transitions Myr^-1^ in Onagraceae, 0.23 transitions Myr^-1^ in Primulaceae, 0.02 transitions Myr^-1^ in *Solanum*, and 0.14 transitions Myr^-1^ in Pooideae. The four phylogenies, with marginal reconstructions of ploidy states at nodes rather than time slices, are depicted in Appendix S1-S4. Pooideae was the clade with the highest number of estimated ploidy transitions with 43 polyploidizations and 53 diploidizations across all times slices; the next largest number of events was in Primulaceae, with 7 polyploidizations and 5 diploidizations. Onagraceae underwent 3 polyploidizations and one diploidization, while we reconstructed 12 polyploidizations and 0 diploidizations in *Solanum*.

Very few events of polyploidization and diploidization were recovered during the M2 and LIG slices, likely due to the small size of those slices (about 100,000 years each). For that reason, our discussion on latitudinal movements in relation to changes in ploidy category is mainly based on results across all time slices.

### Relationships between ploidy shifts and latitudinal movements

ANOVA results indicate that every clade except Pooideae showed significantly higher present-day absolute latitudes in polyploids relative to diploids. This means that we were able to recover the LPG for most clades, namely, polyploids generally were located at higher latitudes than diploids (Fig. 3). However, phylogenetic ANOVA revealed no significant differences between absolute latitudes in diploids and polyploid species in any clade (phylogenetic ANOVA: Onagraceae *F*=27.77, *p*=0.257; Primulaceae *F*=7.168, *p*=0.124; *Solanum F*=8.574, *p*=0.171; Pooideae *F*=0.143, *p*=0.856. This suggests that the observed differences between ploidy states among species were not more different than expected by chance alone, and likely as a consequence of ploidy being less labile than changes in latitude.

**Figure 3.**
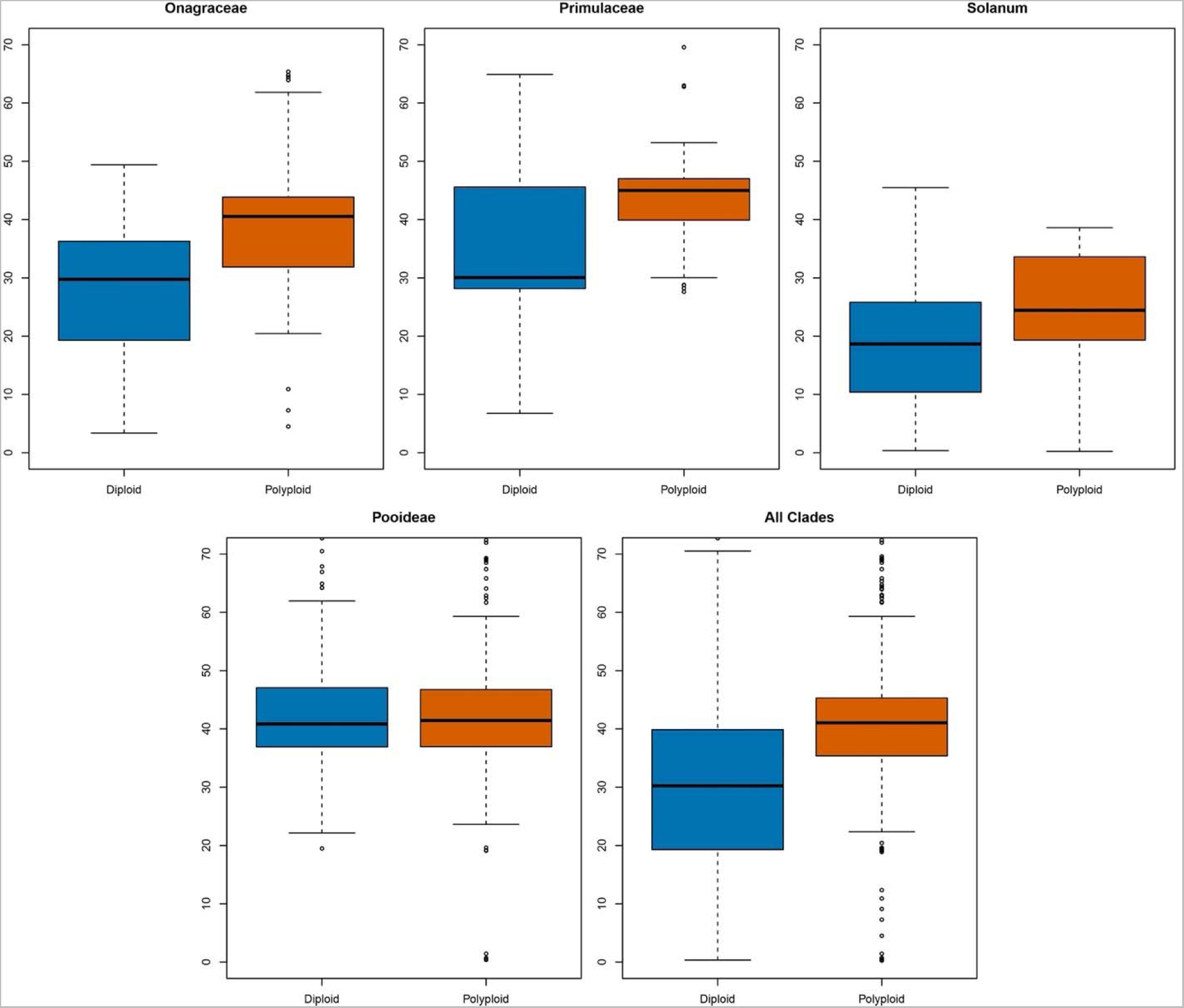
Boxplot showing present-day absolute latitudes of all plants in our dataset by ploidy and by clade.

When we correlated inferred ploidy shifts at particular time slices with estimated starting latitudes for that time slice, we found mixed support for the “centers of origin” hypotheses across clades. Across all clades, phylogenetic paired ANOVA detected almost no significant differences between the starting latitudes of lineages that polyploidize in a given time slice and lineages in the other ploidy status categories (Fig. 4a,b,c,e). The only significant comparison was between species that polyploidized vs. stayed diploid in Pooideae, where those that polyploidize exhibited significative higher starting latitudes than those that stay diploid (*F*=14.109, *p*=0.011; Fig. 4d). Since corHMM reconstructed only one diploidization event in Onagraceae, and zero in *Solanum*, comparisons between the polyploidized and diploidized ploidy status groups could not be conducted in these clades.

**Figure 4.**
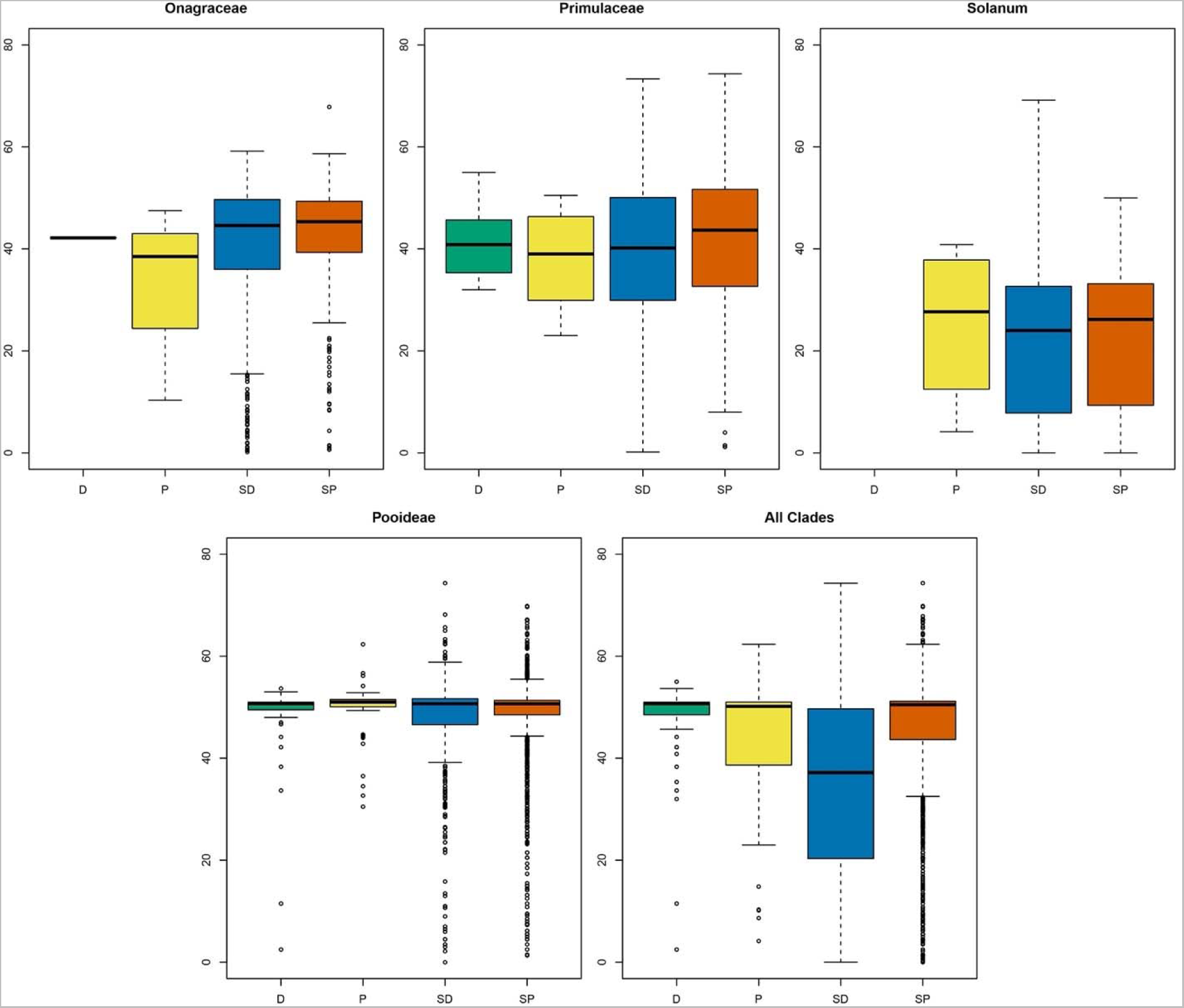
Boxplots comparing the starting absolute latitudes across each of the four ploidy groups, across all time slices, divided by clade. Ploidy status categories from left to right in each plot are: diploidized (“D”), polyploidized (“P”), stayed diploid (“SD”), and stayed polyploid (“SP”).

Regarding support for “centers of arrival,” phylogenetic ANOVA did not support a significant difference in latitudinal movement between lineages that diploidized as opposed to polyploidized in any clade nor across all clades (Fig. 5). A binomial sign test detected marginally significant directional movement only in species that polyploidized in Primulaceae (*p*=0.0625), which moved on average 9.11 degrees latitude antiequatorially, as expected under the “center of arrival” hypothesis. However, a Wilcoxon signed-rank test, which accounts for both direction and magnitude of movement, was not significant for species that either diploidized (*p*=0.8125) or polyploidized (*p*=0.375) in the family. The only significant Wilcoxon test was found for species that polyploidized in Pooideae, both with (*p*=0.0358) and without (*p*=0.019) phylogenetic correction. In this case, lineages that polyploidized tended to move equatorially, rather than antiequatorially. Kolmogorov-Smirnov tests for differences between the changes in median latitudes of species that polyploidized vs. diploidized were all insignificant. Species that polyploidize did show noticeable spikes in northward movement relative to other groups in some clades and time slices (Supplementary Material), but these findings were countered by the mostly insignificant Wilcoxon and Kolmogorov-Smirnov test results.

**Figure 5.**
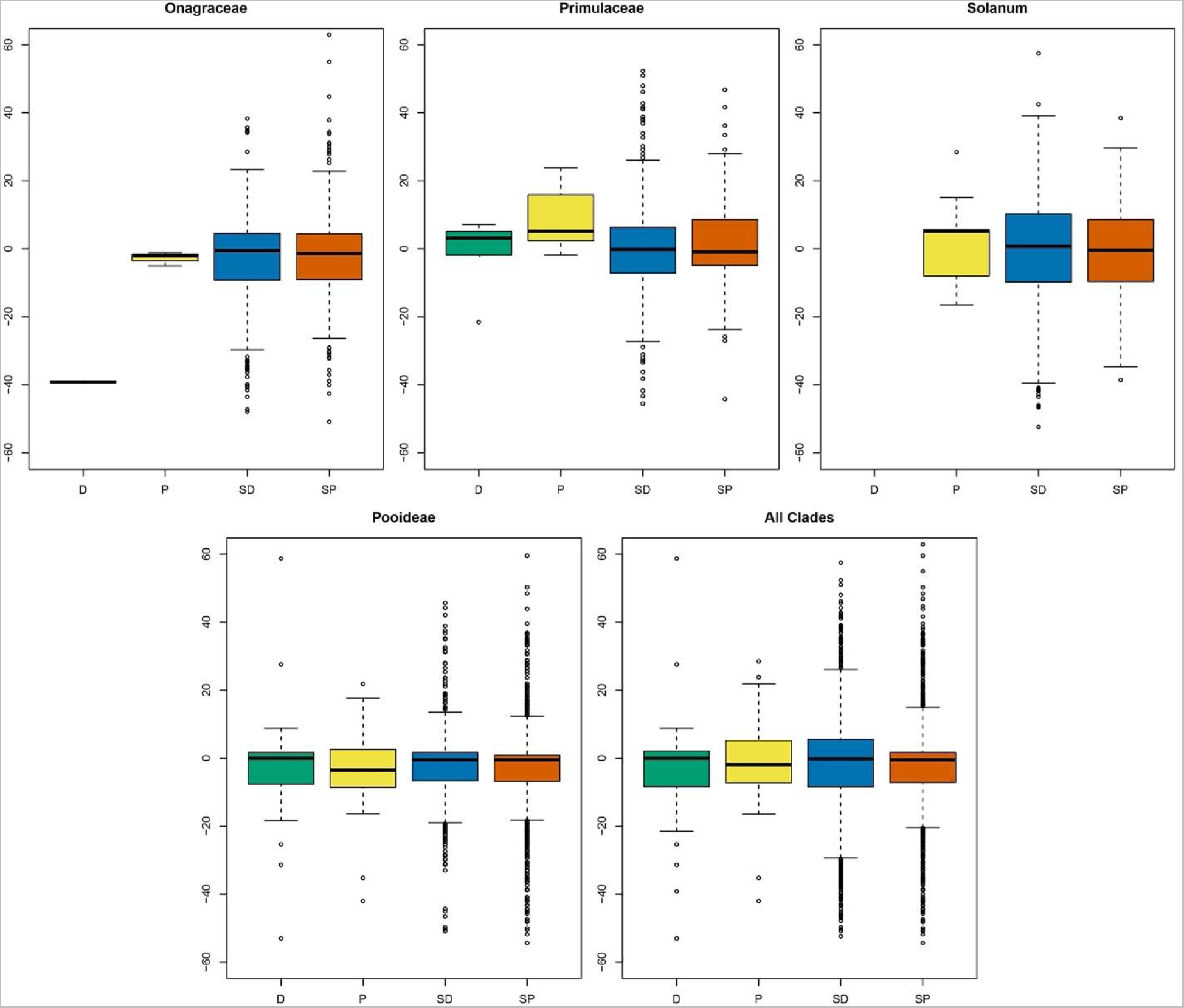
Boxplots comparing the change in median latitude across clades, averaged across all time slices and separated by ploidy status category. Ploidy status categories from left to right in each plot are: diploidized (“D”), polyploidized (“P”), stayed diploid (“SD”), and stayed polyploid (“SP”).

## DISCUSSION

### Causes of the LPG

Our study aimed to determine whether the LPG was better explained by greater rates of origination in or movement into poleward environments by polyploid flowering plants relative to diploid ones. At this point, given the low amount of support for either hypothesis, we do not favor one hypothesis over the other when it comes to general patterns. However, we note that a lack of support for the LPG once relationships between modern day latitude and ploidy take the phylogeny into account (Fig. 3). That could mean that many of the polyploid species in higher latitudes are closely related, which could also be interpreted as support for the “centers of origin” hypothesis.

The general lack of support for either greater rates of movement or origination at high latitudes relative to diploids may accord with a third hypothesis to explain the LPG: that of higher latitude environments being “centers of survival” for polyploid plants. In this scenario, polyploids do not originate or move to poleward latitudes at higher rates relative to diploids. Rather, polyploids and diploids originate at the same rates in high latitude environments, but diploids go extinct more frequently than polyploids (see Stebbins, 1984). In this case, the harsh environmental conditions hypothesized under “centers of origin” to create the conditions for higher rates of unreduced gamete formation, and thus polyploidization, instead filter out diploids in favor of polyploids, perhaps due to polyploidy conferring beneficial traits to tolerate abiotic stresses (Tossi et al., 2022). Our current analysis is unable to detect this possible pattern because we do not examine diversification rates, and strong support for “centers of survival” would involve finding higher rates of extinction in diploids at high latitudes relative to polyploids. We hope that future work will address this and other possible mechanisms for the LPG.

We hasten to acknowledge that the study conducted here is very much a preliminary one. Despite our unclear results, we hope that phylogenetic-informed ecological niche modeling will continue to be used to study both the LPG and other biogeographic patterns. Such methods would be improved by the introduction of more sophisticated ancestral state reconstruction. In machuruku, ancestral characters are estimated assuming a simple Brownian motion model of evolution, and parameters underlying the evolutionary model are not free for the user to adjust or conduct model selection procedures. Fortunately, new work is being conducted studying whether bioclimatic variables are correlated with diversification rate changes (Zhang et al., 2021) and allowing for selective models of climatic evolution like Ornstein-Uhlenbeck models with hidden states (Boyko et al., 2023).

### Clade-specific patterns

We were surprised to find little difference between movements in polyploidized vs. diploidized species in *Solanum*, as this is the only one of our groups that is distributed primarily in the southern rather than northern hemisphere (Olmstead and Palmer, 1997). It is possible that their Andean center of richness causes species to move elevationally rather than latitudinally, though there is a noticeable spike in antiequatorial movement in lineages that polyploidized during the MIS19 (Supplementary Material). Their Andean distribution may also explain the equatorial movement seen in *Solanum* during the mPWP. In the temperate clades, latitudinal differences are difficult to decipher, possibly due to the narrow and biased GBIF ranges centered on Europe (see Beck et al., 2014). The largest group with the most reconstructed ploidy shifts, Pooideae, showed the most significant results by far, with the most tests showing significant differences in latitudinal movement among groups, though movement is most clearly observed in species that diploidized during the LIG (Appendix S6). It is possible that the other, smaller clades with few reconstructed ploidy shifts leave us with little statistical power to detect associations between ploidy and latitudinal movement.

Alternatively, it is possible that the biogeographic patterns displayed by species in each ploidy status category, which compose the LPG, arose prior to the time scale we studied, such as during one of the Pliocene glaciation events in northern latitudes prior to the M2 (De Schepper et al., 2014). In this scenario, species may not exhibit significant movement in the present day or recent geologic past due to niches already being filled in polar environments. Additionally, there is the possibility that the LPG is created via polyploid formation due to secondary contacts of previously isolated populations confined to glacial refugia (Stebbins, 1984, 1985). Though very similar to the “centers of origin” hypothesis, testing this hypothesis would require comparing polyploid frequencies in deglaciated areas to non-deglaciated areas rather than a simple latitudinal comparison. If this is the case, it would explain the lack of movement mostly observed in temperate clades, which possess ranges that overlap with potential glacial refugia (Comes and Kadereit, 1998).

While we did find support for antiequatorial movement in Pooideae in species that polyploidized relative to those that diploidized, and the opposite pattern in Primulaceae, these findings may be better explained by methodological limitations rather than clade-specific traits. While rates of diploidization vary across species (Li et al., 2021), it is likely that full genome reorganization requires much more time than was included in in the 3.3 million years for which we possessed paleoclimatic data (Lynch and Conery, 2003; Landis et al., 2018). Future studies may benefit from examining longer time scales than we considered here.

### Caveats

Our study is not without important caveats. First, ploidy levels of tropically distributed plant species remain largely uncharacterized relative to those with temperate distributions (Husband et al., 2013; Vasconcelos, 2023). This pattern is reflected in a large European bias in the distributions of plants included in this study. Additionally, our interpretations of how ploidy changes relate to subsequent latitudinal movements are limited by the available resolution of paleoclimatic data through time. For example, species that exhibit small amounts of latitudinal change after ploidy change may have transitioned soon before the end of the time slice, and movement in the subsequent time slice that may be caused by the ploidy change would not be detected by our methods. In other words, it is possible that latitudinal movement may occur after a “lag” (Schranz et al., 2012). While the lag hypothesis focuses on gaps between polyploidy and diversification, if lags are often required for the “success” of polyploids, this may also explain delayed ecological shifts or phenotypic shifts that enable range expansion and alteration.

Finally, the unevenness of our historical data makes it difficult to solidly connect ploidy shifts to subsequent latitudinal changes. Our time slices range widely in size: the gap between the M2 and the mPWP is smaller than 100,000 years, while our largest gap between the mPWP and MIS19 is almost 2.5 million years. The large number of ploidy shifts detected between the mPWP and the MIS19 could be attributed to climatic changes, as mean annual temperature declines during this period (Lisiecki and Raymo, 2005), or to the relatively long period between these time slices. Additionally, while the inclusion of phylogeny in reconstructing ancestral ranges will, in theory, produce better predictions, estimates can be spurious in cases where closely related species on the phylogeny exhibit widely disjunctive ranges. In the case of Pooideae, species in the genus *Aciachne* were reconstructed to have a very high median latitude around 50 degrees north in the MIS19 and prior. However, all three species of the genus included in our study have present-day median latitudes around -10 degrees south of the equator, making such large shifts suspect. This is likely driven by the biogeographic influence of closely related genera like *Oryzopsis*, in which all three of the species included in our dataset retain median latitudes around the range of 40 to 50 degrees north from the M2 to the present, and *Piptocheatium*, in which species ranges vary widely. As examples, *P. lasianthum* currently occurs in southern Brazil and northeast Argentina, while *P. avenaceum* occurs from Mexico to southeast Canada (POWO, 2023).

## CONCLUSIONS

Our first examination of the historical causes of the latitudinal polyploidy gradient found clade-specific differences in support for whether the pattern is driven more by polyploid origination at higher latitudes or polyploid movement to higher latitudes. When comparing the median latitudes and latitudinal movement across species that stayed polyploid, stayed diploid, polyploidized, and diploidized in individual time slices, we found significant differences in our largest clade, Pooideae. We also found significant differences in starting latitudes across clades, though the latitudinal relationship between species that polyploidized vs. diploidized varied.

While we were able to detect the LPG in differences in median latitudes occupied by species that stay polyploid as opposed to stay diploid, we likely lack sufficient data to detect differences between species that polyploidize as opposed to diploidize. We hope that further studies using similar methods will re-investigate this question with different, larger clades.

## ACKNOWLEDGMENTS

We thank members of the Beaulieu lab for helpful discussions about the ideas presented here.

## AUTHOR CONTRIBUTIONS

E.R.H. and J.M.B. designed the study. T.V. collected the data. E.R.H., T.V., and J.D.B. analyzed the data and interpreted the results. E.R.H., T.V., J.D.B., and J.M.B. wrote and edited the manuscript.

## DATA AVAILABILITY STATEMENT

All scripts and data generated by this study will be made available through the Dryad Digital Repository upon acceptance.

## Notes

### Competing Interest Statement

The authors have declared no competing interest.

## LITERATURE CITED

1. Akaike, H. 1974. A new look at statistical model identification. IEEE Transactions on Automatic Control AU-19: 716–722.

2. Barringer, B.C. 2007. Polyploidy and self-fertilization in flowering plants. American Journal of Botany 94: 1527–1533.

3. Beaulieu, J.M., B.C. O’Meara, and M.J. Donoghue. 2013. Identifying hidden rate changes in the evolution of a binary morphological character: the evolution of plant habit in campanulid angiosperms. Systematic Biology 62: 725–737.

4. Beck, J., M. Böller, A. Erhardt, and W. Schwanghart. 2014. Spatial bias in the GBIF database and its effect on modeling species’ geographic distributions. Ecological Informatics 19: 10–15.

5. Bierzychudek, P. 1985. Patterns in plant parthenogenesis. Experientia 41: 1255–1264.

6. Brochmann, C., A.K. Brysting, I.G. Alsos, L. Borgen, H.H. Grundt, A.-C. Sheen, and R. Elven. 2004. Polyploidy in arctic plants. Biological Journal of the Linnean Society 82: 521– 536.

7. Boyko, J.D., and J.M. Beaulieu. 2021. Generalized hidden Markov models for phylogenetic comparative datasets. Methods in Ecology and Evolution 12: 468–478.

8. Boyko, J.D., E.R. Hagen, J.M. Beaulieu, and T. Vasconcelos. 2023. The evolutionary responses of life-history strategies to climatic variability in flowering plants. New Phytologist.

9. Brown, J.L., D.J. Hill, A.M. Dolan, A.C. Carnaval, and A.M. Haywood. 2018. PaleoClim, high spatial resolution paleoclimate surfaces for global land areas. Scientific Data 5: 180254.

10. Burnham, K.P., and D.R. Anderson. 2002. Model selection and inference: a practical information-theoretic approach, 2nd ed. Springer-Verlag: New York, New York, USA.

11. Caetano, D.S., B.C. O’Meara, and J.M. Beaulieu. 2018. Hidden state models improve state dependent diversification approaches, including biogeographical models. Evolution 72: 2308–2324.

12. Comes, H.P., and J.W. Kadereit. 1998. The effect of Quaternary climatic changes on plant distribution and evolution. Trends in Plant Science 3: 432–438.

13. Conover, W.J. 1971. Practical nonparametric statistics. John Wiley & Sons: New York, New York, USA.

14. De Schepper, S., P.L. Gibbard, U. Salzmann, and J. Ehlers. 2014. A global synthesis of the marine and terrestrial evidence for glaciation during the Pliocene Epoch. Earth-Science Reviews 135: 83–102.

15. De Storme, N., and D. Geelen. 2014. The impact of environmental stress on male reproductive development in plants: biological processes and molecular mechanisms. *Plant*, Cell & Environment 37: 1–18.

16. De Vos, J.M., R.O. Wüest, and E. Conti. 2014. Small and ugly? Phylogenetic analyses of the “selfing syndrome” reveal complex evolutionary fates of monomorphic primrose flowers. Evolution 68: 1042–1057.

17. Felsenstein, J. 1981. A likelihood approach to character weighting and what it tells us about parsimony and compatibility. Biological Journal of the Linnean Society 16: 183–196.

18. Freyman, W.A., and S. Höhna. 2019. Stochastic character mapping of state-dependent diversification reveals the tempo of evolutionary decline in self-compatible Onagraceae lineages. Systematic Biology 68: 505–519.

19. Goldberg, E.E., L.T. Lancaster, and R.H. Ree. 2011. Phylogenetic inference of reciprocal effects between geographic range evolution and diversification. Systematic Biology 60: 451– 465.

20. Guillory, W.X., and J.L. Brown. 2021. A new method for integrating ecological niche modeling with phylogenetics to estimate ancestral distributions. Systematic Biology 70: 1033–1045.

21. Husband, B.C., S.J. Baldwin, and J. Suda. 2013. The incidence of polyploidy in natural plant populations: major patterns and evolutionary processes. In Leitch, I.J., J. Greilhuber, J. Dolezel, and J. Wendel [eds.], Plant genome diversity, vol. 2, 255–276. Springer, Vienna, Austria.

22. Jiao, Y, N.J. Wickett, S. Ayyampalayam, A.S. Chanderbali, L. Landherr, P.E. Ralph, L.P. Tomsho, et al. 2011. Ancestral polyploidy in seed plants and angiosperms. Nature 473: 97– 100.

23. Landis, J.B., D.E. Soltis, Z. Li, H.E. Marx, M.S. Barker, D.C. Tank, and P.S. Soltis. 2018. Impact of wholeLgenome duplication events on diversification rates in angiosperms. American Journal of Botany 105: 348L363.

24. Leitch, A.R., and I.J. Leitch. 2008. Genomic plasticity and the diversity of polyploid plants. Science 320: 481–483.

25. Levin, D.A. 1983. Polyploidy and novelty in flowering plants. The American Naturalist 122: 1–25.

26. Li, Z., M.T.W. McKibben, G.S. Finch, P.D. Blischak, B.L. Sutherland, and M.S. Barker. 2021. Patterns and processes of diploidization in land plants. Annual Review of Plant Biology 72: 387–410.

27. Lisiecki, L.E., and M.E. Raymo. 2005. A PlioceneLPleistocene stack of 57 globally distributed benthic δ18O records. Paleoceanography 20: PA1003.

28. Lohaus, R., and Y. Van de Peer. 2016. Of dups and dinos: evolution at the K/Pg boundary. Current Opinion in Plant Biology 30: 62–69.

29. Löve, A., and D. Löve. 1943. The significance of differences in the distribution of diploids and polyploids. Hereditas 29: 145–163.

30. Löve, A., and D. Löve. 1949. The geobotanical significance of polyploidy. I. Polyploidy and latitude. In Goldschmidt, R.B. [ed.], Portugaliae Acta Biologica, Série A special volume, p. 273–352. Instituto Botânico de Lisboa, Lisbon, Portugal.

31. Lynch, M., and J.S. Conery. 2003. The evolutionary demography of duplicate genes. In Meyer, A., and Y. Van de Peer [eds.], Genome evolution: gene and genome duplications and the origin of novel gene functions, 35–44. Springer, Dordrecht.

32. Ma, Z., B. Sandel, and J.-C. Svenning. 2016. Phylogenetic assemblage structure of North American trees is more strongly shaped by glacial-interglacial climate variability in gymnosperms than in angiosperms. Ecology and Evolution 6: 3092–3106.

33. Mandáková, T., and M.A. Lysak. 2018. Post-polyploid diploidization and diversification through dysploid changes. Current Opinion in Plant Biology 42: 55–65.

34. Marks, R.A., S. Hotaling, P.B. Frandsen, and R. VanBuren. 2021. Representation and participation across 20 years of plant genome sequencing. Nature Plants 7: 1571–1578.

35. Matzke, N.J. 2014. BioGeoBEARS: BioGeography with Bayesian (and Likelihood) Evolutionary Analysis in R Scripts.

36. Mayrose, I., S.H. Zhan, C.J. Rothfels, K. Magnuson-Ford, M.S. Barker, L.H. Rieseberg, and S.P. Otto. 2011. Recently formed polyploid plants diversify at lower rates. Science 333: 1257.

37. McElwain, J.C., and M. Steinthorsdottir. 2017. Paleoecology, ploidy, paleoatmospheric composition, and developmental biology: a review of the multiple uses of fossil stomata. Plant Physiology 174: 650–664.

38. Meyers, L.A., and D.A. Levin. 2006. On the abundance of polyploids in flowering plants. Evolution 60: 1198–1206.

39. Mudelsee, M., and Raymo, M.E. 2005. Slow dynamics of the Northern Hemisphere glaciation. Paleoceanography 20: PA4022.

40. Normand, S., R.E. Ricklefs, F. Skov, J. Bladt, O. Tackenberg, and J.-C. Svenning. 2011. Postglacial migration supplements climate in determining plant species ranges in Europe. Proceedings of the Royal Society B 278: 3644–3653.

41. Olmstead, R.G., and J.D. Palmer. 1997. Implications for the phylogeny, classification, and biogeography of *Solanum* from cpDNA restriction site variation. Systematic Botany 22: 19– 29.

42. POWO. 2023. Plants of the World Online. Royal Botanic Gardens, Kew. http://www.plantsoftheworldonline.org/

43. Price, T.D., A. Qvarnström, and D.E. Irwin. 2003. The role of phenotypic plasticity in driving genetic evolution. Proceedings of the Royal Society B 270: 1433–1440.

44. R Core Team. 2022. R: A language and environment for statistical computing. R Foundation for Statistical Computing: Vienna, Austria. https://www.R-project.org/.

45. Ramsey, J., and D.W. Schemske. 2002. Neopolyploidy in flowering plants. Annual Review of Ecology and Systematics 33: 589–639.

46. Raunkiaer, C. 1934. The life forms of plants and statistical plant geography; being the collected papers of C. Raunkiaer. Clarendon Press, Oxford, UK.

47. Ree, R.H., and S.A. Smith. 2008. Maximum likelihood inference of geographic range evolution by dispersal, local extinction, and cladogenesis. Systematic Biology 57: 4–14.

48. Revell, L.J. 2012. phytools: an R package for phylogenetic comparative biology (and other things). Methods in Ecology and Evolution 3: 217–223.

49. Rice, A., P. Šmarda, M. Novosolov, M. Drori, L. Glick, N. Sabath, S. Meiri, et al. 2019. The global biogeography of polyploid plants. Nature Ecology & Evolution 3: 265–273.

50. Robertson, K., E.E. Goldberg, and B. Igić. 2011. Comparative evidence for the correlated evolution of polyploidy and self-compatibility in Solanaceae. Evolution 65: 139–155

51. Särkinen, T., L. Bohs, R.G. Olmstead, and S. Knapp. 2013. A phylogenetic framework for evolutionary study of the nightshades (Solanaceae): a dated 1000-tip tree. BMC Evolutionary Biology 13: 1–15.

52. Schranz, M.E., S. Mohammadin, and P.P. Edger. 2012. Ancient whole genome duplications, novelty and diversification: the WGD Radiation Lag-Time Model. Current Opinion in Plant Biology 15: 147–153.

53. Segraves, K.A. 2017. The effects of genome duplications in a community context. New Phytologist 215: 57–69.

54. Smirnov, N.V. 1939. On the estimation of the discrepancy between empirical curves of distribution for two independent samples. Moscow University Mathematics Bulletin 2: 3–26.

55. Soltis, D.E., M.C. Segovia-Salcedo, I. Jordon-Thaden, L. Majure, N.M. Miles, E.V. Mavrodiev, W. Mei, et al. 2014. Are polyploids really evolutionary dead-ends (again)? A critical reappraisal of Mayrose et al. (2011). New Phytologist 202: 1105–1117.

56. Soreng, R.J., P.M. Peterson, K. Romaschenko, G. Davidse, J.K. Teisher, L.G. Clark, P. Barberá, et al. 2017. A worldwide phylogenetic classification of the Poaceae (Gramineae) II: an update and a comparison of two 2015 classifications. Journal of Systematics and Evolution 55: 259–290.

57. Spriggs, E.L., P.-A. Christin, and E.J. Edwards. 2014. C4 photosynthesis promoted species diversification during the Miocene grassland expansion. PloS ONE 9: e97722.

58. Stebbins, G.L. 1950. Variation and evolution in plants. Columbia University Press, New York, New York, USA.

59. Stebbins, G.L. 1971. Chromosomal evolution in higher plants. Addison-Wesley, London, UK.

60. Stebbins, G.L. 1984. Polyploidy and the distribution of the arctic-alpine flora: new evidence and a new approach. Botanica Helvetica 94: 1–13.

61. Stebbins, G.L. 1985. Polyploidy, hybridization, and the invasion of new habitats. Annals of the Missouri Botanical Garden 72: 824–832.

62. Sundaram, M., and A.B. Leslie. 2021. The influence of climate and palaeoclimate on distributions of global conifer clades depends on geographical range size. Journal of Biogeography 48: 2286–2297.

63. Svenning, J.-C., and F. Skov. 2007. Could the tree diversity pattern in Europe be generated by postglacial dispersal limitation? Ecology Letters 10: 453–460.

64. Tossi, V.E., L.J. Martinez Tosar, L.E. Laino, J. Iannicelli, J.J. Regalado, A.S. Escandón, I. Baroli, H.F. Causin, and S.I. Pitta-Álvarez. 2022. Impact of polyploidy on plant tolerance to abiotic and biotic stresses. Frontiers in Plant Science 13: 869423.

65. Van Drunen, W.E., and B.C. Husband. 2019. Evolutionary associations between polyploidy, clonal reproduction, and perenniality in the angiosperms. New Phytologist 224: 1266–1277.

66. Vasconcelos, T. 2023. A trait-based approach to determining principles of plant biogeography. American Journal of Botany 110: e16127.

67. Wilcoxon, F. 1945. Individual comparisons by ranking methods. Biometrics Bulletin 1: 80–83.

68. Xu, Z., and L. Chang. 2017. Primulaceae. In Xu, Z., and L. Chang [eds.], Identification and control of common weeds, vol. 3, 51–81. Springer, Singapore.

69. Yang, Z. 2006. Computational molecular evolution. Oxford University Press, London, UK.

70. Zenil-Ferguson, R., J.G. Burleigh, W.A. Freyman, B. Igić, I. Mayrose, and E.E. Goldberg. 2019. Interaction among ploidy, breeding system and lineage diversification. New Phytologist 224: 1252–1265.

71. Zhang, X., J.B. Landis, Y. Sun, H. Zhang, N. Lin, T. Kuang, X. Huang, T. Deng, H. Wang, and H. Sun. 2021. Macroevolutionary pattern of Saussurea (Asteraceae) provides insights into the drivers of radiating diversification. Proceedings of the Royal Society B 288: 20211575.

