## Supporting Information for "Historical causes for the greater proportion of polyploid plants in higher latitudes"

### *American Journal of Botany* Supporting Information

Article acceptance date: NA

The following Supporting Information is available for this article:

**Appendix S1.** Phylogeny of Onagraceae, pruned from the phylogeny by Freyman and Höhna (2019). Dotted lines near the tips represent the four time slices for which we possessed paleoclimatic data. Time in millions of years is shown at the top. Marginal reconstructions of ploidy states with traditional corHMM are shown at the nodes. Blue colors represent diploids, red colors represent polyploids, and gray colors represent tips with no ploidy data.

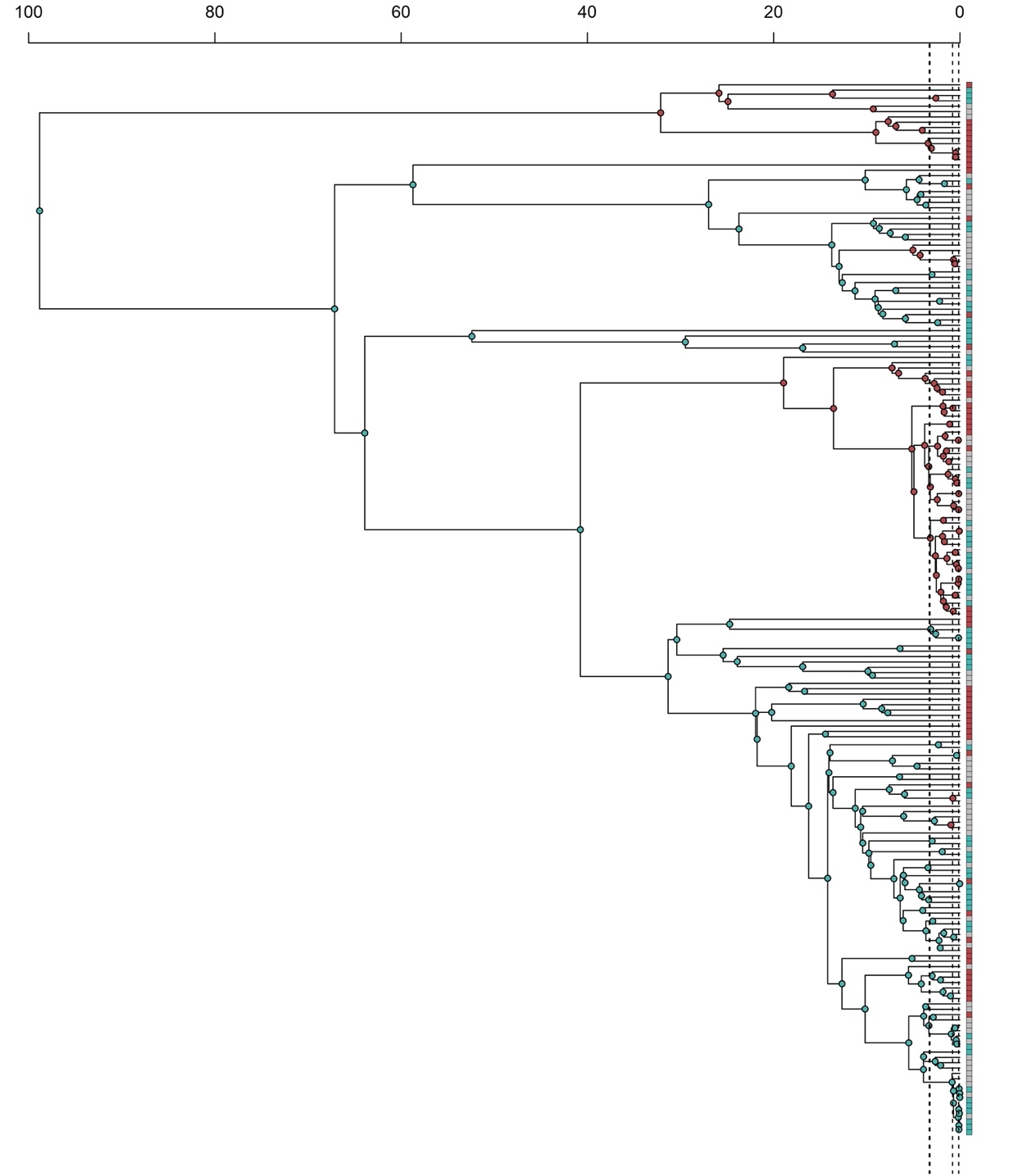

**Appendix S2.** Phylogeny of Pooideae, pruned from the phylogeny of Poaceae by Spriggs et al. (2014). Dotted lines near the tips represent the four time slices for which we possessed paleoclimatic data. Time in millions of years is shown at the top. Marginal reconstructions of ploidy states with traditional corHMM are shown at the nodes. Blue colors represent diploids, red colors represent polyploids, and gray colors represent tips with no ploidy data.

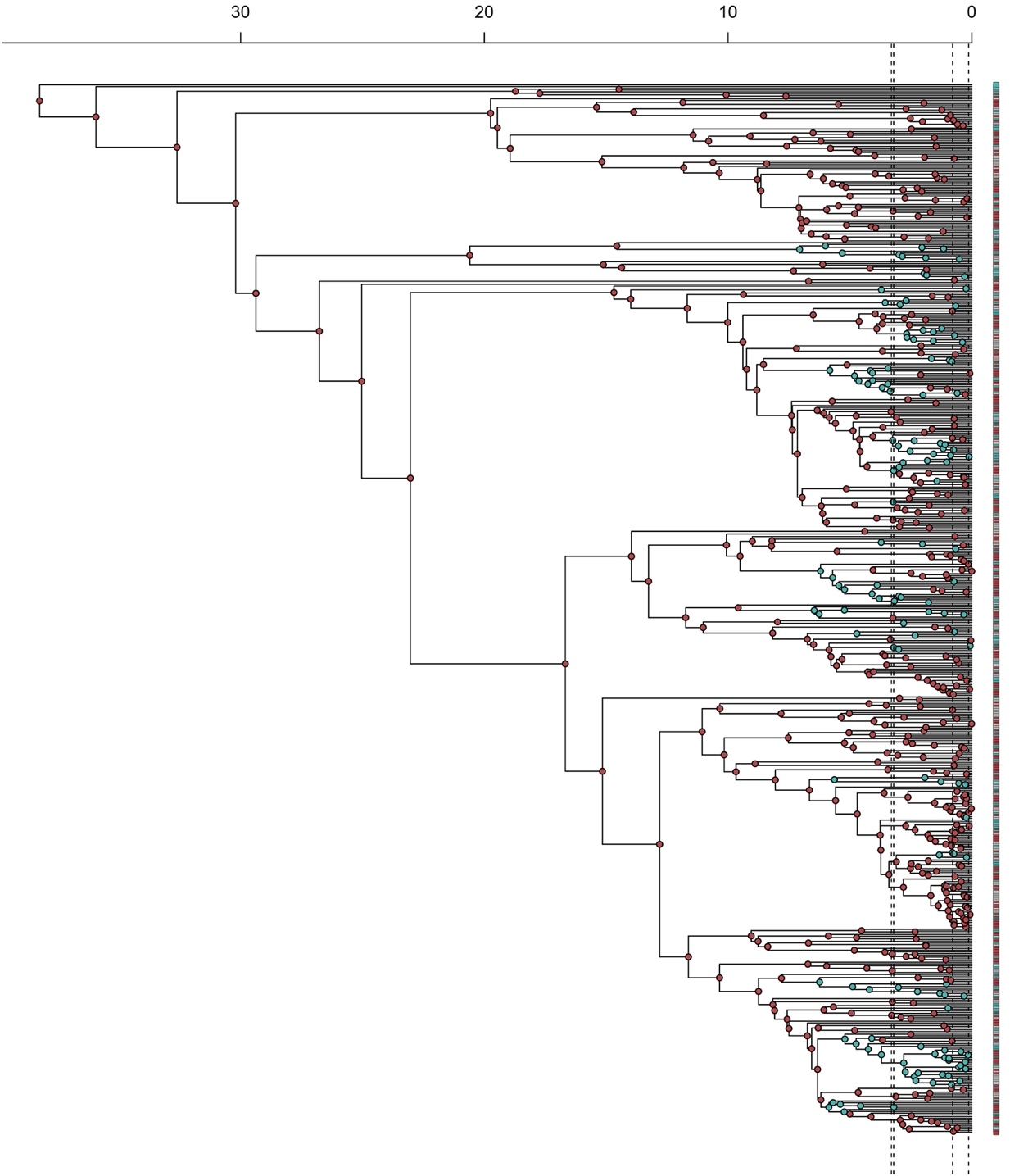

**Appendix S3.** Phylogeny of Primulaceae, pruned from the phylogeny by De Vos et al. (2014). Dotted lines near the tips represent the four time slices for which we possessed paleoclimatic data. Time in millions of years is shown at the top. Marginal reconstructions of ploidy states with traditional corHMM are shown at the nodes. Blue colors represent diploids, red colors represent polyploids, and gray colors represent tips with no ploidy data.

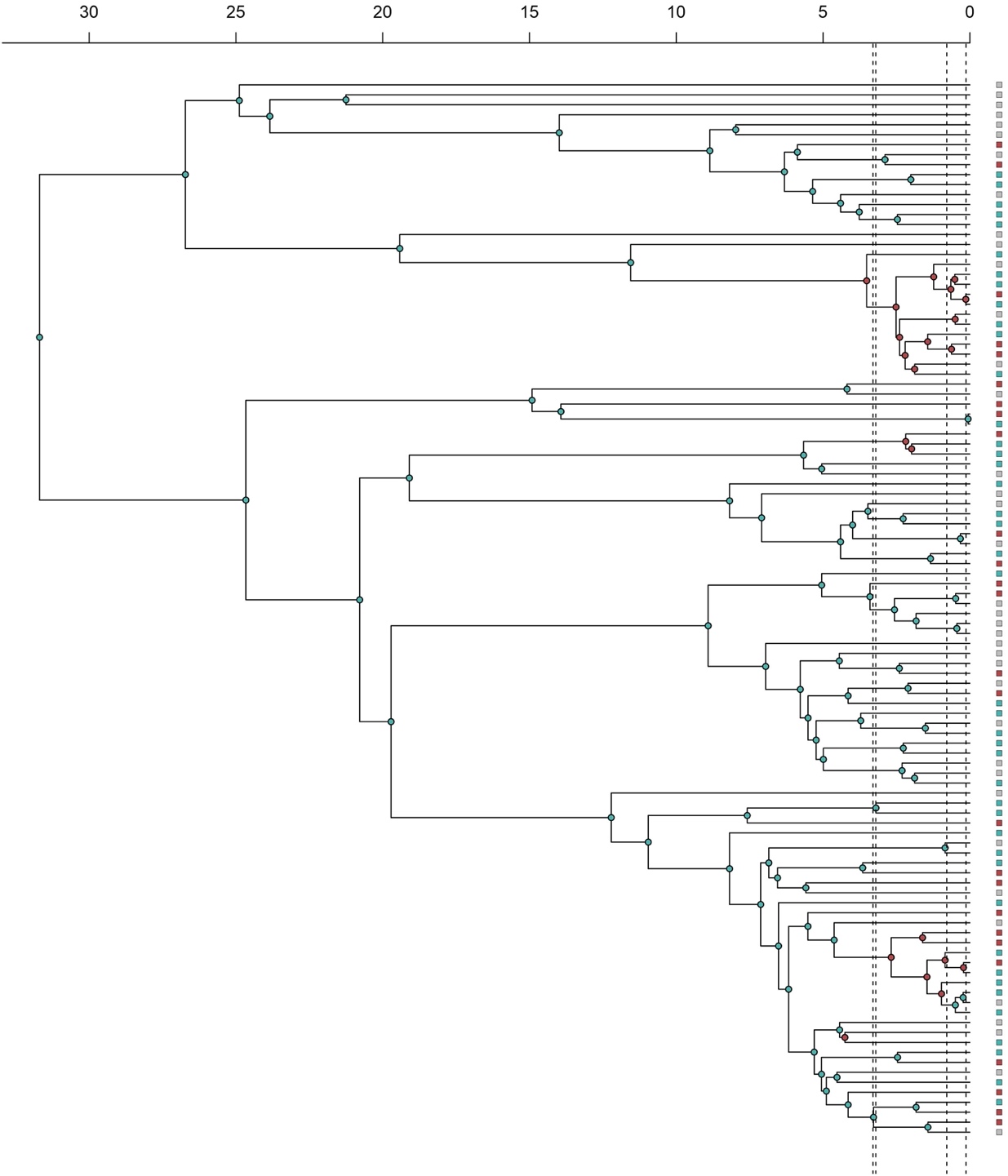

**Appendix S4.** Phylogeny of *Solanum* (Solanaceae), pruned from the phylogeny of Solanaceae by Särkinen et al. (2013). Dotted lines near the tips represent the four time slices for which we possessed paleoclimatic data. Time in millions of years is shown at the top. Marginal reconstructions of ploidy states with traditional corHMM are shown at the nodes. Blue colors represent diploids, red colors represent polyploids, and gray colors represent tips with no ploidy data.

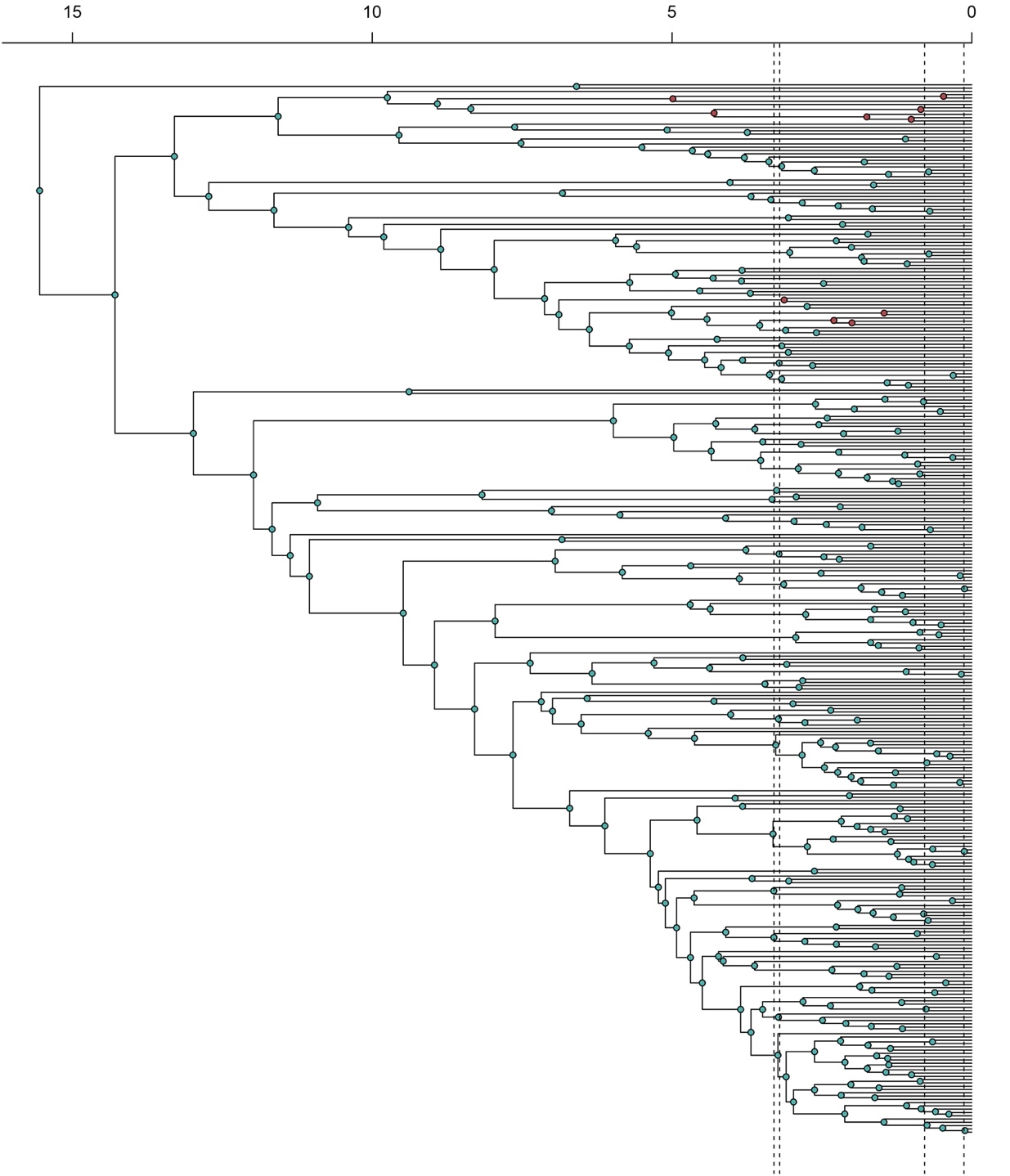

**Appendix S5.** Boxplots comparing the starting absolute latitudes within individual time slices and clades, divided by ploidy status group. Ploidy status categories from left to right in each plot are: diploidized (“D”), polyploidized (“P”), stayed diploid (“SD”), and stayed polyploid (“SP”). Each column corresponds to a time slice (labeled at top with the beginning slice) and each row corresponds to a clade (with movement across all clades for each time slice in the bottom row).

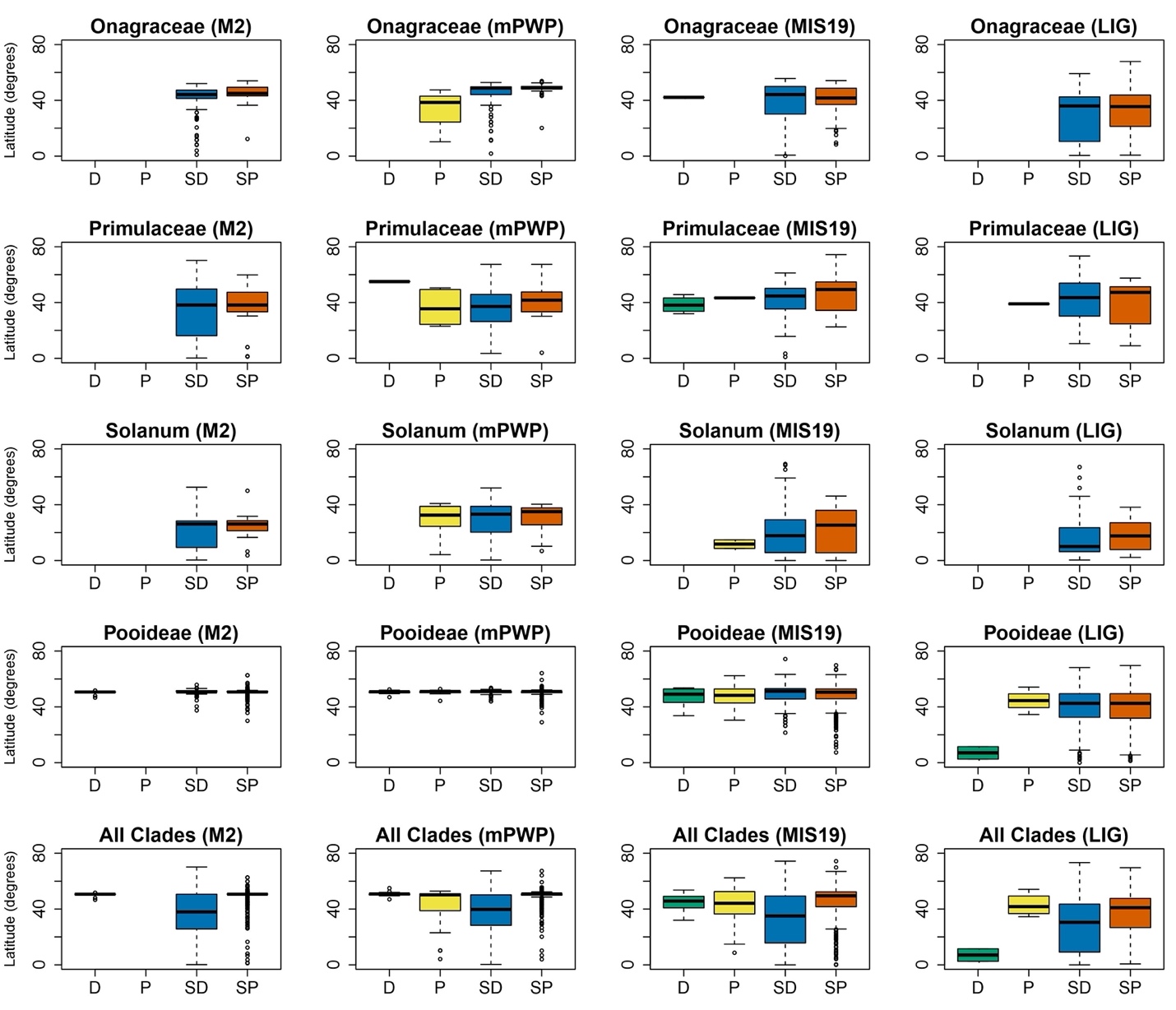

**Appendix S6.** Boxplots comparing the change in median latitude within individual time slices and clades, divided by ploidy status group. Ploidy status categories from left to right in each plot are: diploidized (“D”), polyploidized (“P”), stayed diploid (“SD”), and stayed polyploid (“SP”). Each column corresponds to a time slice (labeled at top with the beginning slice) and each row corresponds to a clade (with movement across all clades for each time slice in the bottom row).

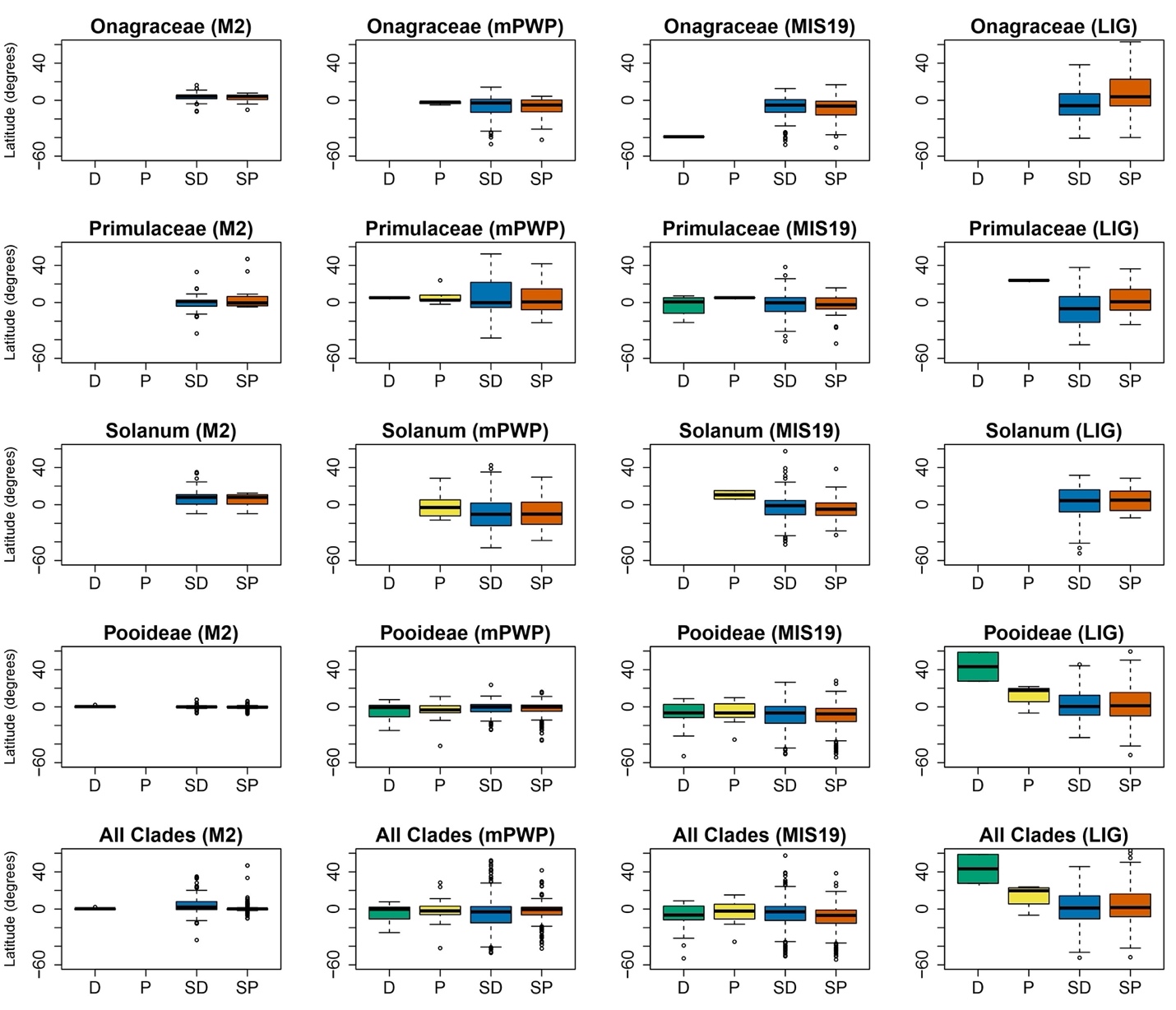
